# Core-periphery structure in mutualistic networks: an epitaph for nestedness?

**DOI:** 10.1101/2020.04.02.021691

**Authors:** Ana M. Martín González, Diego P. Vázquez, Rodrigo Ramos-Jiliberto, Sang Hoon Lee, Vincent Miele

**Affiliations:** Pacific Ecoinformatics and Computational Ecology Lab, Berkeley, CA, USA; Center for Macroecology, Evolution and Climate, University of Copenhagen, Copenhagen, Denmark; Argentine Institute for Dryland Research, CONICET, CC 507, 5500 Mendoza, Argentina; Faculty of Exact and Natural Sciences, National University of Cuyo, Padre Jorge Contreras 1300, M5502JMA Mendoza, Argentina; GEMA Center for Genomics, Ecology & Environment, Faculty of Interdisciplinary Studies, Universidad Mayor, Camino La Pirámide 5750 Huechuraba, Santiago, Chile; Department of Liberal Arts, Gyeongnam National University of Science and Technology, Jinju, Korea; Université de Lyon; Université Lyon 1; CNRS, UMR5558, Laboratoire de Biométrie et Biologie Evolutive, F-69622 Villeurbanne, France

**Keywords:** core-periphery, ecological networks, mesoscale, nestedness, stochastic block models topology

## Abstract

The calculation of nestedness has become a routine analysis in the study of ecological networks, as it is commonly associated with community resilience, robustness and species persistence. While meaningful in species distributional patterns, for an interaction matrix to be nested, specialist species must interact with ordered subsets of subsequently more generalized species — not just with a lower number of species. However, after reviewing 419 papers on mutualistic networks published since nestedness was introduced for the study of species interactions in 2003, we have found that only two theoretical studies considered explicitly ordered subsets. Instead, most studies interpret nestedness as a core of densely connected generalist species, surrounded by a periphery of specialist species attached to this core — a so-called core-periphery structure. Such a topological feature is generally perceived as a core-periphery structure in network science. Here, we argue that the concept of core-periphery may be more relevant for studies on mutualistic networks than the concept of nestedness, as ecologists are usually not interested in exploring in detail the ordered subsets that characterize nestedness but instead use nestedness to describe a topology with a core of densely linked generalist species surrounded by a sparsely linked periphery of specialists. To illustrate our arguments and the quantification of core-periphery structures, we calculate core-periphery and nestedness in a large publicly available dataset of mutualistic networks. We also describe the calculation of core-periphery structures, its relationship with nestedness, and provide the code inside the R package *econetwork* for its calculation in mutualistic networks. We hope that our review will help ecologists to move beyond nestedness towards a more explicit representation of the structure of ecological networks.

## Nestedness in ecology

Some of the key challenges in ecology concern describing the patterns — and understanding the processes — determining the spatio-temporal distribution of species and their interactions, within and among ecosystems (May, 1999; Sutherland et al., 2013). For instance, in the past years, many studies have aimed at identifying community assembly rules through the characterization of networks of biotic interactions (e.g., Bascompte et al., 2003; Olesen et al., 2008; Ulrich & Gotelli, 2010; Jonhson et al., 2013). By examining biotic interactions at the community level, research moved beyond the study of pairwise interactions or interactions among small subsets of species, which offered novel insights into the ecological and evolutionary dynamics of entire communities of interacting species.

A routine metric for the characterization of the structural organization of ecological systems is nestedness: the analysis of nested subsets of interaction among species. Nestedness was first proposed in biogeography for the study of species’ beta-diversity (Patterson, 1986; Atmar & Patterson, 1993; Ulrich et al., 2009). The spatial distribution of species sets is typically analyzed as species by site matrices, with species as rows, sites as columns, and cells indicating the presence/absence or the abundance of each species in each site (Atmar & Patterson, 1993; Ulrich et al., 2009; Ulrich & Gotelli, 2010). If species distributions are nested, species-poor sites are proper subsets of species-rich sites systematically (Patterson & Atmar, 1986; Wright & Reeves, 1992; Wright et al., 1997; Figure 1a). In a species-site matrix where rows and columns are ordered from left to right and top to bottom by a decreasing number of interactions, filled cells will be concentrated at the upper left corner of the matrix. Nestedness is interpreted as the result of ecological processes determining species range dynamics, such as orderly sequences of colonization/extinction dynamics, species/area relationships, environmental filtering or habitat gradients (Patterson & Atmar, 1986; Ulrich et al., 2009; Ulrich & Almeida-Neto, 2012).

**Figure 1.**
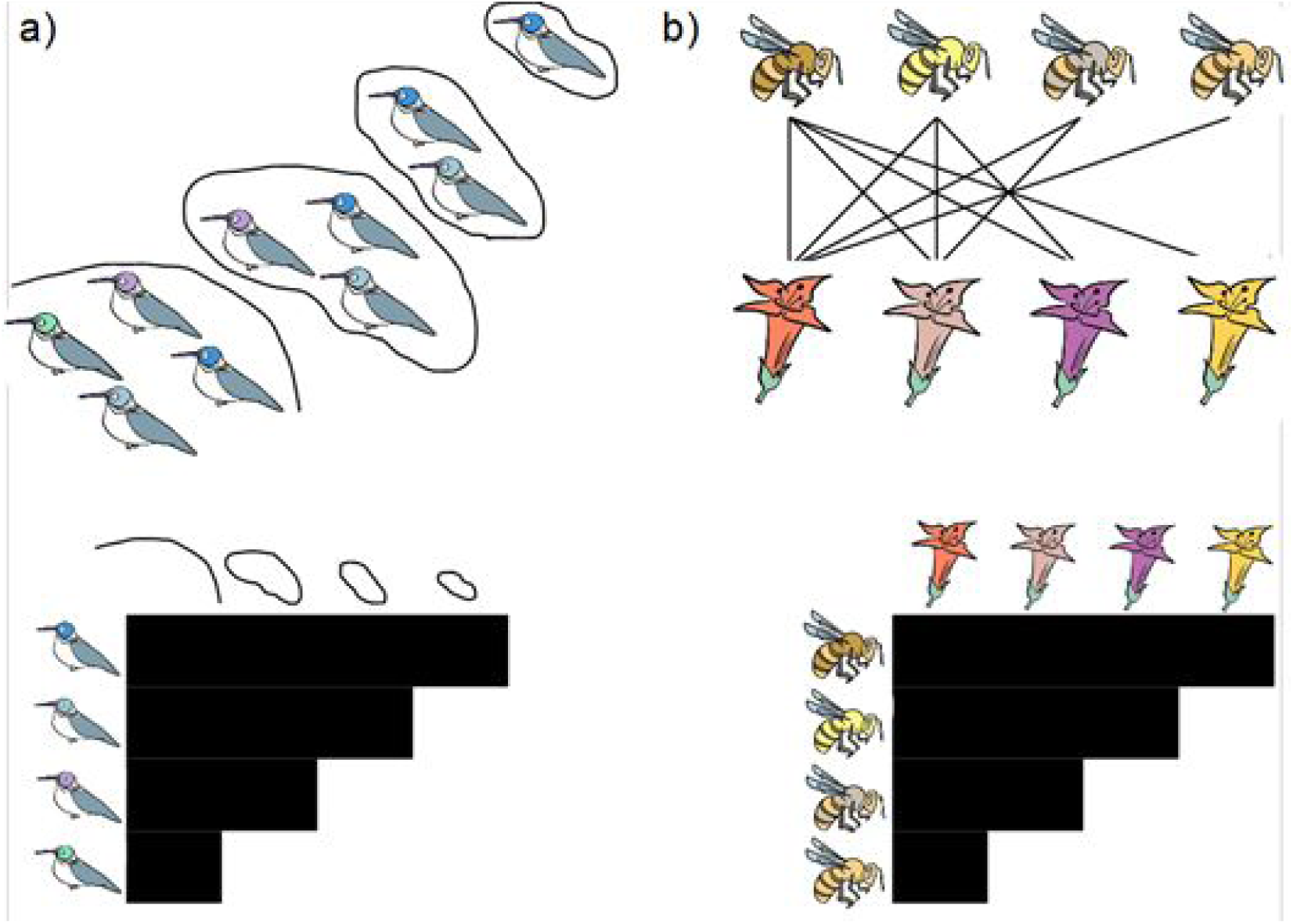
Diagram describing the concept of nestedness using examples from spatial distributions of bird species on islands of decreasing size and distance from the source (a) and interaction patterns between plant and pollinator species (b). On top, representation of the birds in each island (a) and network diagram showing connections (links) between bees and their visited flowers. In the bottom, the same situations in a matrix format, where filled cells represent spatial distributions or interactions between species. We kindly thank @inmunosoni for the drawings.

The initial studies considering nestedness in plant-animal mutualistic networks by Bascompte et al. (2003), Dupont et al. (2003) and Ollerton et al. (2003) introduced the concept and calculation of nestedness for the study of entire communities of mutualistic interaction, quickly becoming a routine analysis in the characterization of mutualistic networks. A search in the Web of Science database with the keywords TOPIC: (nestedness AND (mutualis* OR frugivor* OR pollinat* OR seed dispers*), and limiting the search to articles published since 2003 in science-related journals resulted in 469 studies, of which 415 examined some type of biotic interaction (Supporting Information 1). As distribution matrices, mutualistic networks are bipartite, i.e., composed of two different groups of entities such as plants and pollinators, or plants and seed dispersers. In a nested matrix with rows and columns ordered by a decreasing number of interactions, interactions will be concentrated on the upper left corner of the matrix. Hence, nestedness measures whether a given species interacts with subsequent subsets of the interaction partners of more generalized species, as *“a set of nested Russian Matryoshka dolls”* (Bascompte, 2010; Fig 1b). In studies on mutualistic interactions, nestedness is regarded as a sign of generalization, as it implies a lack of interactions among specialized species, which would appear at the bottom right corner of the ordered matrix; and of specialization reciprocity, as specialists interact with more generalized species. As a result, in a nested network there is a high cohesiveness, through a highly connected, although undefined, network core, and a high asymmetry, with specialist species interacting mostly with generalists at the core (Bascompte et al. 2003). Thus, the use of nestedness to characterize mutualistic networks represented a conceptual breakthrough, confirming the change of paradigm from a highly reciprocally specialized conception on mutualistic interactions to one of widespread asymmetric specialization, in which many specialized species interact with a core of highly generalized species (Bascompte et al., 2003; Dupont et al., 2003; Ollerton et al., 2003; Vázquez & Aizen, 2004).

Since then, a vast number of studies routinely calculate nestedness to describe networks of mutualistic interactions. Moreover, as most communities show significantly higher nestedness than randomized networks (e.g. Bascompte et al., 2003), numerous studies suggest that a nested organization of the interactions at the community level increase biodiversity and the robustness of the interacting community, and provide more resilience to habitat loss (Fortuna & Bascompte, 2006; Bastolla et al., 2009; Thébault & Fontaine, 2010); but see James et al. (2012) and Valdovinos et al. (2016)).

## What nestedness measures…

The concept of nestedness, as defined in the early biogeographic studies, implies a consistent ordering in hierarchical subsets, with species-poor sites containing subsets of the species present in progressively richer sites (Fig 1a). The various nestedness metrics available focus on these orderings, although in different ways (Ulrich et al., 2009; Fig 2). Podani & Schmera (2011) attempted some redefinition of nestedness and presented alternative metrics which, unlike most commonly used metrics, do not depend on matrix ordering. Their approach was contested by a review on nestedness by Ulrich & Almeida-Neto (2012) stressing the importance of using metrics that focus specifically on the row/column gradients in order to identify the ecological drivers responsible for the nested subsets of interactions. Nevertheless, it is still unclear which outcomes of nestedness analysis give the most relevant insights for understanding the structure of mutualistic networks. This is a serious problem, as there are several metrics and software platforms currently available leading to different interpretations about the degree and significance of nestedness (Figure 2, SM2; Csermely et al., 2013; Ulrich & Almeida-Neto, 2012).

**Figure 2.**
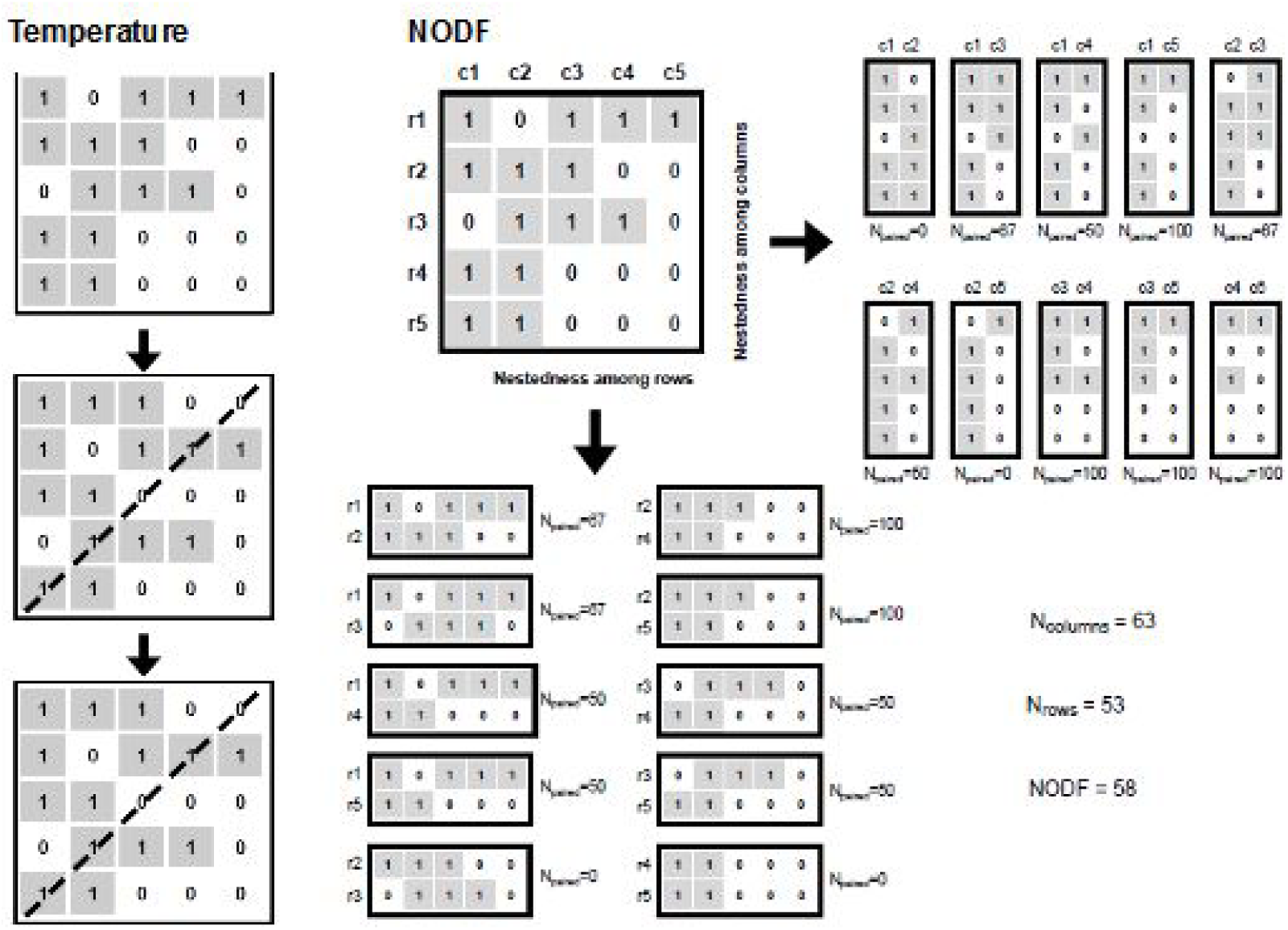
Illustration depicting how the two nestedness metrics most used in ecological interaction networks are calculated. The top row contains the original matrix, and the bottom row details about the nestedness computation. Notice that both metrics are based on hierarchical nested subsets of the observed interactions. A) Nestedness temperature *T* (Atmar & Patterson 1995). The diagonal represents the isocline of perfect order, so that in a perfectly nested matrix, all 1’s will be above the isocline, and all 0’s below. Temperature measures all unexpected absences above and presences below the isocline, weighting them by the square Euclidean distance to the isocline. B) Nestedness based on Overlap and Decreasing Fill *NODF.* NODF quantifies the overlaps for all combinations of column and row pairs with *decreasing marginal totals*, divided by the total number of combinations of column and row pairs (Almeida-Neto et al. 2008). Figure adapted from Almeida-Neto et al. 2008.

The first metric used to calculate nestedness in interaction networks was *Temperature*, a measure of the deviation from a theoretically perfectly nested matrix (Fig. 2; Atmar & Patterson, 1993). Yet, Temperature (and other temperature-based metrics, such as BINMATNEST, as much as other popular metrics such as discrepancy or C), have important limitations, as they are not scrupulous enough with the ordering of rows and columns, measure only the deviation from the matrix’s own maximum nestedness instead of from an independent benchmark, or are non-convergent computationally, resulting in inconsistencies when calculating nestedness in matrices of varying size and connectance (Guimarães & Guimarães, 2006; Rodriguez-Girones & Santamaria, 2006; Almeida-Neto et al., 2008). To overcome these challenges, Almeida-Neto and colleagues introduced in 2008 “*nestedness based on overlap and decreasing fill*” (NODF), which has become the most widely used metrics in mutualistic networks. The NODF metric is regarded to represent more accurately the original concept of nestedness by using the average values of paired rows and columns to measure directly whether specialized species interact with proper subsets of the partners of more generalized species in an ordered way (Almeida-Neto et al., 2008; Ulrich et al., 2009; Csermely et al., 2013). Importantly, however, the calculation of NODF implies that any deviation from such progressive filling has a strong impact on the resulting value (Figure 2). This feature of NODF could be problematic, as mutualistic networks may have missing interaction data throughout the entire matrix, including among core, generalists and *a priori* thoroughly observed species (Bosch et al., 2009).

## …and what we think it measures in interaction network data

Although the measurement of ordered subsets is fundamental for the definition, calculation and interpretation of the ecological drivers of nested patterns (Csermely et al., 2013; Ulrich & Almeida-Neto, 2012), in the numerous publications assessing nestedness in mutualistic interaction networks there is an astounding lack of investigation and discussion about whether observed interactions in natural communities are in fact ordered in nested subsets, let alone what mechanism drives that pattern (Supplementary Information). To our knowledge, only two recent studies by Burkle and Runyon (2019) and Losapio et al. (2019), examine some level of detail into the hierarchical subsetting of interactions. In the former, species’ nestedness ranks are associated with their VOCs emissions along a flowering season, whereas in the latter the authors included plant-plant facilitation and competition interactions in a plant-pollinator interaction network, examining in detail interaction rewiring between species and its effect on nestedness. There are other several studies that incorporate nested subsets of interactions (115 studies, 28%), but they do so exclusively from a theoretical perspective (either by introducing new metrics to overcome biases and limitations in the calculation of nestedness, or describe theoretical models of community build-up; Supplementary Information). In all the other published studies on mutualistic networks, the interpretation of nestedness is reduced to a description of the existence of a generalist core of species hypothesized to drive network dynamics, and a periphery of specialized species attached to this core (e.g. Bascompte et al., 2003).

## Why a core-periphery structure?

Network mesoscale structures derive from node degree distributions, which in mutualistic networks are highly skewed, with few highly generalized species and a long tail of specialists (Jordano et al., 2003; Jonhson et al., 2013; Sajjad et al., 2017). Numerous studies suggest that observed degree distributions result from neutral processes, such as highly skewed abundances or sampling-related issues, which ultimately result in large differences in species’ detection probabilities (Ollerton et al., 2003; Vázquez & Aizen, 2004; Vázquez et al., 2005, 2009; Kallimanis et al., 2009; Araujo et al., 2010). For instance, current practices involve the quantification of network properties on data that is aggregated temporally, disregarding temporal dynamics despite these communities show high turnovers of species and interactions, with a highly skewed distribution of species’ phenophases, and a network structure that evolves seasonally (Ollerton et al., 2003; Vázquez et al., 2005, 2009; Olesen et al., 2008; Petanidou et al., 2008; Kallimanis et al., 2009; Araujo et al., 2010; Zhang et al., 2011; Falcão et al., 2016; Sajjad et al., 2017; Kantsa et al., 2018). In addition to skewed degree distributions, mutualistic networks tend to have a high degree of disassortativity, that is, specialist species tend to interact with generalists (Vázquez & Aizen, 2004; Jonhson et al., 2013). The ecological explanation for disassortativity lies in a theoretically higher reliability of generalists, together with higher temporal overlaps of species and a higher detection probability of interactions involving generalist/abundant species (e.g. Martín González et al., 2012). The combination of highly skewed degree distributions and some degree of disassortativity results in a community-wide interaction pattern where generalist species are part of a relatively dense core with specialists forming a periphery attached to this core. For that reason, various studies suggest that the pervasiveness of the nested pattern in mutualistic networks is a mere by-product of such community assembly rules, and not at emergent feature of species interactions as initially proposed (Ollerton et al., 2003; Vázquez et al., 2005, 2009; Araujo et al., 2010; Kallimanis et al., 2009; Jonhson et al., 2013; Valverde et al., 2018; Payrató-Borrás et al., 2019). For instance, the fact that reshuffling interactions within fixed core-periphery blocks results in almost the original nestedness value (Fig. 5d), suggests that the main driver of observed nestedness is the highly skewed link distribution of species.

## Core-periphery as an alternative to nestedness to describe mutualistic networks

A network displaying a core-periphery structure is composed of a core (a subgroup of densely connected nodes) surrounded by a periphery of sparsely connected nodes attached to it (Everett & Borgatti, 1999; Rombach et al., 2012; Lee, 2016; Miele et al., 2020). This is how the majority of studies on mutualistic interaction networks interpret nestedness. However, as explained above, it is not exactly what nestedness measures because nestedness metrics consider, in various forms, the presence of hierarchical subsets of interactions. Therefore, we suggest examining the core-periphery structures instead, and to measure nestedness only when we are interested in exploring whether interactions are ordered in hierarchical subsets. Although the ecological meaning of weighted nestedness is debated (see for instance recent discussion^1^), the ecological interpretation of computing core-periphery in a weighted network is arguably more straightforward—stronger interactions are expected to occur between generalist species at the network core (Miele et al., 2020).

In a matrix displaying a core-periphery structure, there is a species ordering (i.e. an ordering in rows and columns) such that interactions are distributed in an “L-shape”. This L-shape is composed by four blocks of varying connectance: block C11 represents the core, includes all of the interactions between core plants and animals, and thus has the highest connectance; blocks C12 and C21 include the interactions between core plants and animals respectively with animals and plants that are at the periphery, and thus have also a high connectance (in general lower than or close to C11); block C22 includes the interactions that occur between peripheral plants and animals, and has the lowest connectance of all blocks (in general close to 0, Figure 3). Therefore, the three blocks form the L-shape (block C11 is the core, and blocks C12 and C21 are the tails of the degree distribution), with a fourth, extremely sparse block C22. Here we propose to rely on statistical inference to weigh the evidence for the existence of such core-periphery structure (in other words, whether the interaction matrix is organized in the above four blocks). We use the framework of “stochastic block models” (SBMs; Leicht & Newman, 2008; Daudin et al., 2008), originally developed in social sciences and recently applied in ecology (Allesina & Pascual, 2009; Martín González et al., 2012; Leger et al., 2015; Kéfi et al., 2016; Miele et al., 2020). An SBM will infer groups of statistically equivalent nodes, i.e. species that tend to be connected in a similar way, handling binary and weighted networks with the appropriate statistical distribution (Bernouilli for binary data, Poisson for integer weights, and Gaussian for float weights; Mariadassou et al, 2010). A statistically-grounded model selection decides the number of groups that best explain network structure (Daudin et al., 2008). Here, we use SBMs with an additional simple trick: we limit the model to infer a maximum of four groups: blocks C11, C12, C21 and C22. However, it is important to note that the model selection procedure can lead to a model with less than 4 blocks. In this study, we apply this model and consider a network has a core-periphery only when 1) it is described by four blocks, and 2) when these blocks have an L-shape, i.e., connectance in C11 is greater than in C12 and C21, whereas connectance in C22 is also lower than in the three other blocks (see Figure 4). For these networks we can further define a “core-peripheriness” value (CPness) that illustrates the strength of the core-periphery partition, as CPness=(E11+E12+E21)/E, where Eij is the number of interactions (edges) or the sum of weights for each block (E*ij* for block *ij*) or for the entire network (E). This approach was implemented using the R package *blockmodels* (Leger et al., 2015) and is now available as part of the R package *econetwork*.

**Figure 3.**
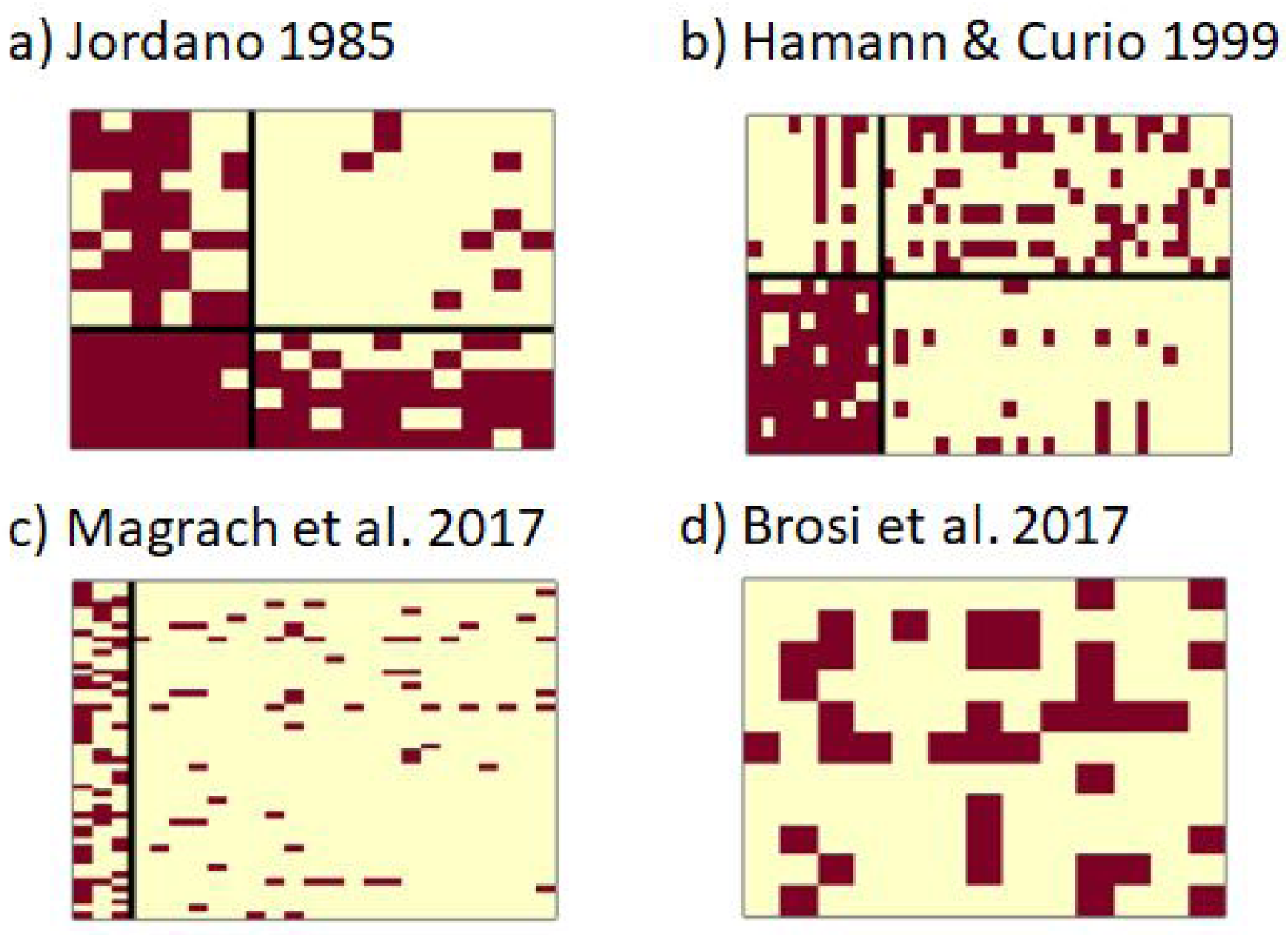
Matrices illustrating four potential block partitions retrieved by the stochastic block model (SBM). Matrices can show a core-periphery structure (Fig. 4a) or not (Fig. 4b-d). Networks that were not core-periphery could be structured into two defined modules, hence not meeting the requirement of a lower connectance in block C22 compared to blocks C12 and C21 (Fig. 4b); be composed of only two blocks, hence only either the plants or the pollinators/seed-dispersers could be divided into core and peripheral species (Fig. 4c); or a total lack of partition (Fig. 4d). Network references (from a-d): Jordano 1985, Hamann & Curio 1999, Magrach et al. 2017 (Sweden lowland community), and Brosi et al. 2017 (Judd Falls TH community).

**Figure 4.**
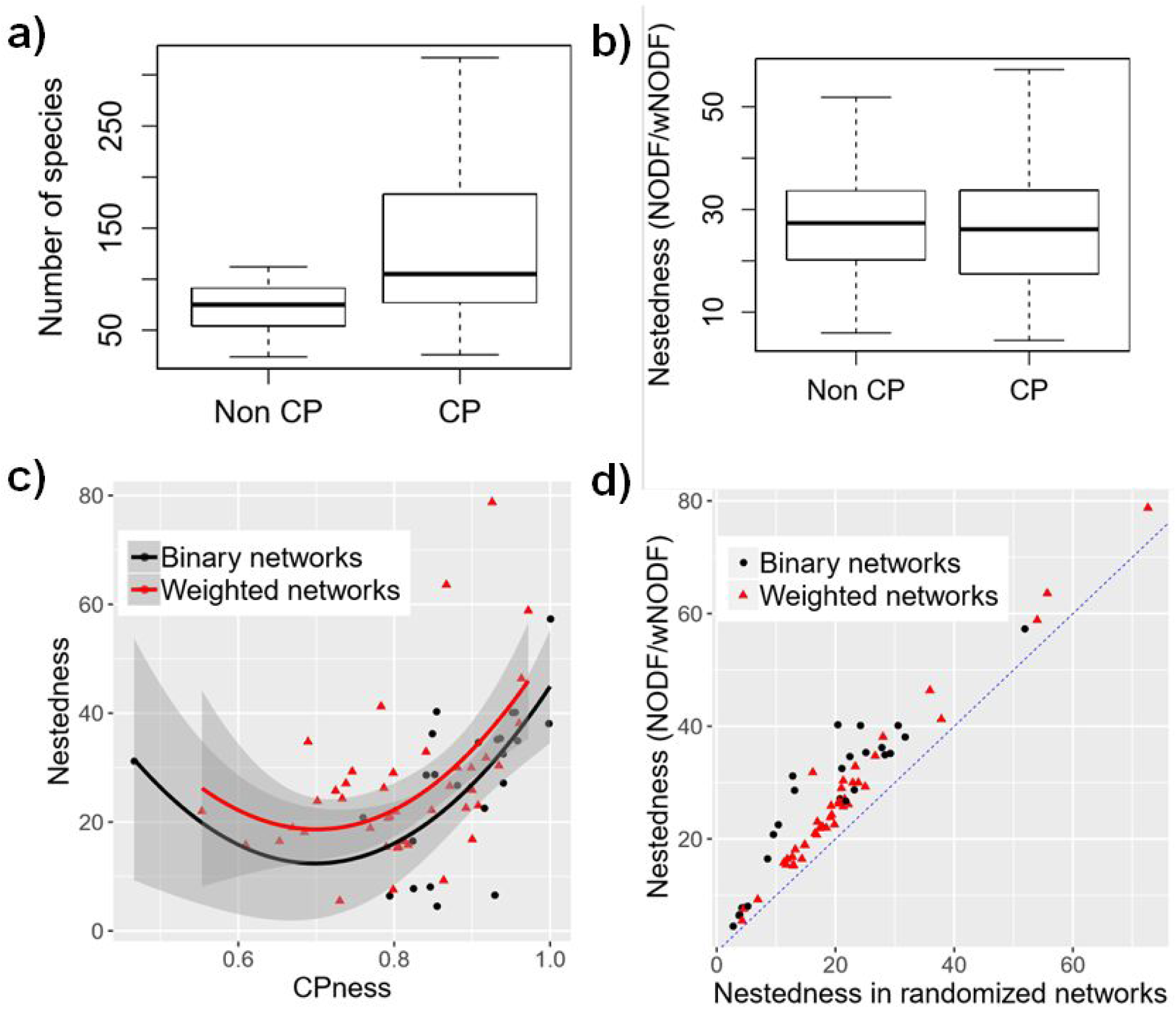
Results from the nestedness and core-periphery analysis. a) Size of networks with and without a core-periphery structure (67 and 44 networks respectively). Notice that networks with a core-periphery structure tend to be larger. b) Distribution of nestedness values in networks with and without a core-periphery structure. Notice that distributions are similar. c) Correlation between Nestedness and core-periphery in binary and weighted networks (43 and 24 networks respectively). d) Correlation between Nestedness calculated on the original matrices and in randomized matrices where interactions have been shuffled within the core-periphery blocks.

To illustrate the relevance of examining the core-periphery of mutualistic networks, we worked with a dataset of 159 binary and weighted mutualistic networks (Michalska-Smith & Allesina, 2019). To ensure that the algorithm has sufficient statistical power, we limited our analysis to networks with at least 10 species per category (e.g. plant and pollinators/seed dispersers, respectively), leaving a total of 111 networks (28 binary, 83 weighted) for the core-periphery analysis. To compare with nestedness, we also computed the NODF2 for binary networks and weighted NODF2 for weighted networks in the package *bipartite* v.2.11 (NODF2 sorts the matrix before calculating NODF, and thus is suited for comparisons across different networks as it is independent of the initial matrix presentation; Dormann et al. 2008).

We detected a core-periphery structure in 67 out of 111 networks examined (60%; Figure 3a). Networks lacking a core-periphery structure were for instance structured into two defined modules, hence not meeting the requirement of a lower connectance in block C22 compared to blocks C12 and C21 (Figure 3b, Figure 4a); composed of two blocks, hence only either the plants or the pollinators/seed-dispersers could be divided into core and peripheral species (Figure 3c); or lacking a partition in any of the groups (Figure 3d). Furthermore, the core-periphery structure tended to be more prevalent in larger networks (Figure 4a). The latter relationship of core-peripheriness and network size could result from the statistical properties of our method, or be a sampling artifact, as sampling completeness may be lower for larger networks (see next section and Jordano, 1987). Interestingly, contrary to our expectations of higher values of nestedness in networks with a significant core-peripheriness compared to networks without a core-periphery, these follow comparable distributions (Figure 4b), suggesting that, despite previous claims (Lee, 2016; Mariani et al. 2019), nestedness and core-periphery structure are not fully equivalent concepts. Nevertheless, they are related: in both binary and weighted networks with a CP structure, the CPness index (that quantifies the number of links inside the L-shape over the total number of links in the network) is positively correlated with network nestedness values (Figure 4c). This relationship is a by-product of the fact that the denser the L-shape, the higher the CPness index, but also the more likely nestedness will be high because of the large number of interactions among generalist plant and animal mutualists (but not much higher than expected on random networks with the same core-periphery structure; see Figure 4d). In these cases, nested sets are found by chance due to the density of links in the L-shape.

## CONCLUSIONS

As our literature review illustrates, nestedness analysis is widespread in the study of plant-animal mutualistic networks. However, as we have argued throughout this manuscript, published studies describe a core-periphery structure, e.g. a topology of a dense core of generalist species and a periphery of specialists attached to the core, without examining whether interactions of specialist species are nested within those of more generalized species. To the best of our knowledge, no study examines whether the progressively ordered subsets associated with any gradient beyond a node’s degree. In fact, a handful of studies have already tried to move away from measuring nestedness, examining core-periphery (or a related aspect) in mutualistic networks. Recently, Miele et al. (2020) used the SBM to examine network structures in a plant-pollinator network over six years, finding a consistent core-periphery structure through seasons and years. Previously, Fang & Huan (2012), Dáttilo et al. (2013) and Partida-Lara et al. (2018) had used *k-cores decomposition* to explore animal-plant mutualistic networks and identify core species. In this study we propose to assess directly the core-periphery structure of mutualistic networks, and have illustrated its usefulness through example calculations on published data. Quantifying a network’s core-periphery allows identifying the cohesive core subgroup, commonly believed to drive overall network stability (e.g. Bascompte et al. 2003, Mariani et al. 2019; Miele et al. 2020). Hence, for detecting species core, we suggest replacing the nestedness analysis with the core-periphery analysis, and encourage future studies to examine the extent to which core-periphery is a general feature of mutualistic networks, and whether this structure and the species composition of the core is consistent across space and time. Furthermore, besides our exploratory work, it would be advantageous to develop new alternative approaches for a more complete and robust description of core-periphery in networks.

## ACKNOWLEDGEMENTS

AMMG was supported through a Marie Skłodowska-Curie Individual Fellowship (H2020-MSCA-IF-2015-704409) and thanks the Danish National Research Foundation for its support of the Center for Macroecology, Evolution and Climate (Grant number DNRF96). DPV was supported through a grant from Argentina’s National Fund for Science and Technology (FONCYT grant no. PICT-2014-3168). RR-J was supported through CONICYT/FONDECYT grant no. 1190173. SHL was supported by the National Research Foundation of Korea (NRF) Grant No. 2018R1C1B5083863. VM was supported by the French National Center for Scientific Research (CNRS) and the French National Research Agency (ANR) grant ANR-18-CE02-0010-01 EcoNet.

## SUPPLEMENTARY MATERIAL

**Supplementary Material 1.** Information on the literature review.

**Supplementary Table 1**. List of articles reviewed. Search query on Web of Science TOPIC=(nestedness AND (mutualis* OR frugivor* OR pollinat* OR seed dispers*), limited to indexed Science Journals for the period 2015-2019.

https://docs.google.com/spreadsheets/d/1b54MrwO7QbeFQcK9CA-sgDkq0F57cgLb4WpX6_y8Y-Y/edit?usp=sharing

**Supplementary Figure 1.**
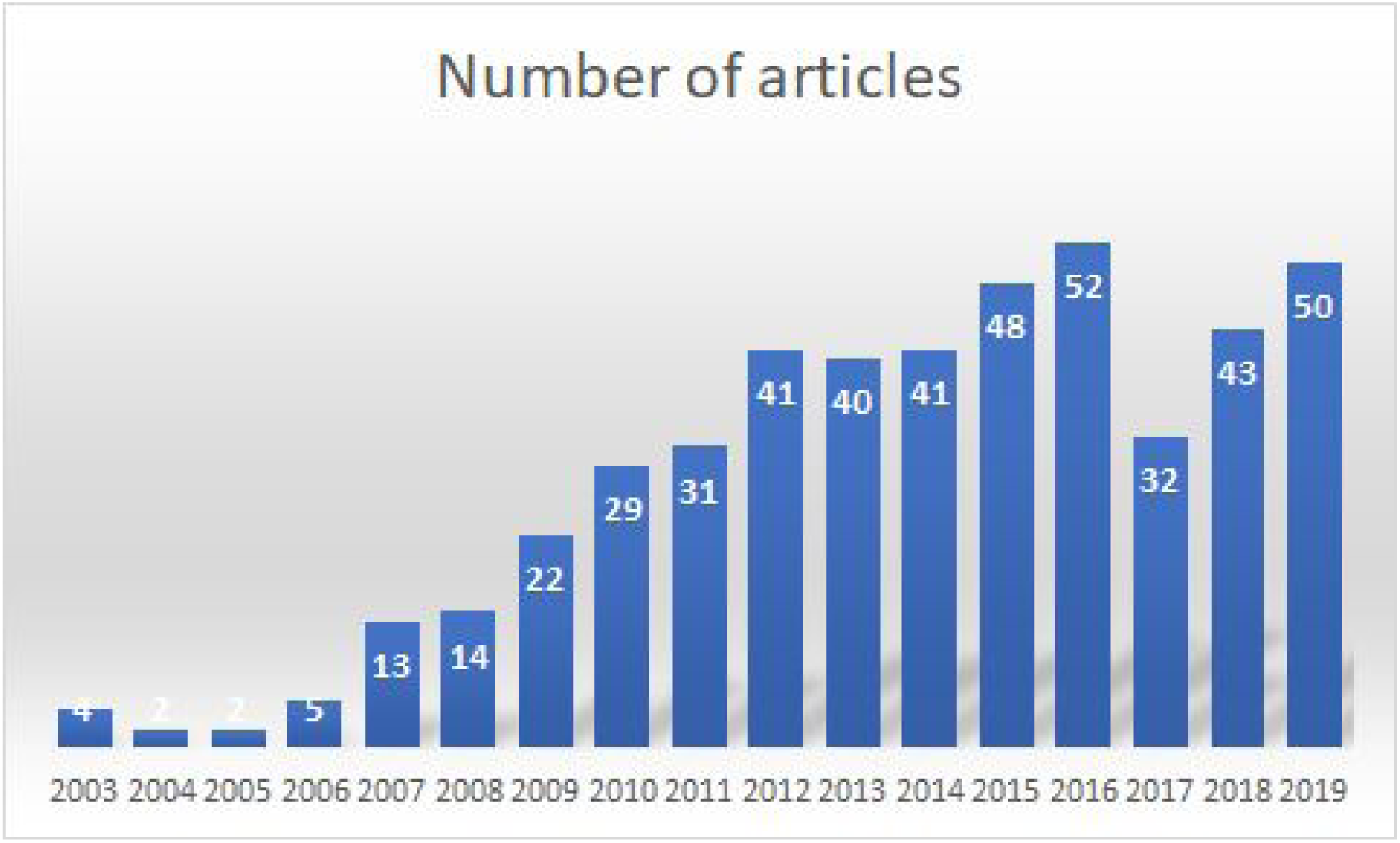
Number of studies published on nestedness per year. Search query on Web of Science TOPIC=(nestedness AND (mutualis* OR frugivor* OR pollinat* OR seed dispers*), limited to indexed Science Journals for the period 2003-2019.

## Footnotes

1 https://jeffollerton.wordpress.com/2019/08/12/weighted-nestedness-and-classical-nestedness-analyses-do-not-measure-the-same-thing-in-species-interaction-networks

